# Repeated evolution of herbicide resistance in *Lolium multiflorum* revealed by haplotype-resolved analysis of acetyl-CoA carboxylase

**DOI:** 10.1101/2023.07.10.548383

**Authors:** Caio A. C. G. Brunharo, Patrick J. Tranel

## Abstract

Herbicide resistance in weeds is one of the greatest challenges in modern food production. The grass species *Lolium multiflorum* is an excellent model species to investigate convergent evolution under similar selection pressure because populations have repeatedly evolved resistance to many herbicides, utilizing a multitude of mechanisms to neutralize herbicide damage. In this work, we investigated the gene that encodes acetyl-CoA carboxylase (ACCase), the target-site of the most successful herbicide group available for grass weed control. We sampled *L. multiflorum* populations from agricultural fields with history of intense herbicide use, and studied their response to three ACCase-inhibiting herbicides under controlled conditions. To elucidate the mechanisms of herbicide resistance and the genetic relationship among sampled populations, we resolved the haplotypes of 97 resistant and susceptible individuals by performing an amplicon-seq analysis using long-read DNA sequencing technologies, focusing on the DNA sequence encoding the carboxyl-transferase domain of ACCase. Our dose-response data indicated the existence of many, often unpredictable, resistance patterns to ACCase-inhibiting herbicides, where populations exhibited as much as 37-fold reduction in herbicide response. The majority of the populations exhibited resistance to all three herbicides studied. Phylogenetic and molecular genetic analyses revealed multiple evolutionary origins of resistance-endowing *ACCase* haplotypes, as well as widespread admixture in the region regardless of cropping system. The amplicons generated were very diverse, with haplotypes exhibiting 26 to 110 polymorphisms. Polymorphisms included insertions and deletions 1-31 bp in length, none of which were associated with the resistance phenotype. We also found evidence that some populations have multiple mechanisms of resistance. Our results highlight the astounding genetic diversity in *L. multiflorum* populations, and the potential for convergent evolution of herbicide resistance across the landscape that challenges weed management and jeopardizes sustainable weed control practices. We provide an in-depth discussion of the evolutionary and practical implications of our results.

## INTRODUCTION

One of the greatest challenges of our generation is to sustainably increase food production to feed a population predicted to grow to 10 billion by 2050 (United Nations, 2017), as well as reduce food insecurity to an estimated 2.4 billion people who currently do not have access to adequate food (FAO, 2021). The sustainable intensification of agriculture is one of the most promising strategies to ensure food security for an exponentially growing world population while protecting natural resources and the environment (Pretty, 2018). Agricultural weeds can interfere with crop growth and development, harbor pests, reduce the marketability of the final product, and can cause the greatest yield loss compared to other pests (Oerke, 2006). Reducing weed interference is, therefore, key for the implementation of sustainable intensification of agriculture. Weed management is primarily performed with herbicides worldwide, and the overreliance on these chemicals has selected for herbicide-resistant populations in many agricultural systems.

The repeated evolution of herbicide resistance in agricultural weeds is a remarkable example of parallel and convergent trait evolution, having been reported in over 1650 weed populations around the world (Heap, 2022). In this work, we use the term “parallel evolution” to refer to resistance conferred by the same genetic modifications at the nucleotide level within a given gene, whereas in “convergent evolution” resistance is achieved by different modifications (for a review of the terms and concepts, see Arendt and Reznick, 2007). Examples of repeated evolution have been extensively observed in *Lolium multiflorum* Lam. populations, having evolved resistance to eight herbicides from different sites of action in 13 countries, often with multiple reports from each country in distinct cropping systems. There are three ecological constraints that reduce the likelihood of resistance evolution. First, after an herbicide application in a field, it is expected that most of the individuals will be killed. The repeated application of herbicides (yearly or multiple times per year) is expected to pose a genetic bottleneck, reducing the standing genetic variation in the population from which weeds can recover [i.e., evolutionary rescue (Carlson et al., 2014)]. The bottleneck can be exacerbated by the use of mixtures of herbicides with different modes of action, a common practice in agriculture. Second, the time between initial herbicide selection pressure and evolution of resistant individuals may be very short, as low as three generations (Busi et al., 2012; Busi & Powles, 2009), and it is unlikely that new mutations arose to confer the resistance phenotype. Third, immigration could contribute to reconstitution of genetic diversity; however, population structure analyses have shown that resistant populations have shared ancestry and limited hybridization (Brunharo & Streisfeld, 2022; Ravet et al., 2021). Recent studies indicated that resistance evolution from standing genetic variation is predominant in the weeds *Amaranthus tuberculatus* (Kreiner et al., 2019) and *Alopecurus myosuroides* (Kersten et al., 2021), although more research is needed to elucidate the evolutionary mechanisms in other species and agricultural contexts.

Herbicide resistance mechanisms in weeds are characterized as either target site or non-target site (reviewed by Gaines et al., 2020). Target-site resistance is conferred by alterations in the enzyme inhibited by the herbicide. These alterations can manifest in the form of mutations in the gene encoding the target site (minimizing or preventing inhibition), or enhanced target enzyme activity by increased gene expression or duplication (Gaines et al., 2010). Non-target-site resistance, on the other hand, involves alterations in plant physiology other than at the herbicide target site. Common physiological modifications observed are enhanced herbicide metabolism (Brunharo et al., 2019) and reduced herbicide translocation (Brunharo & Hanson, 2017). The genetic mechanisms conferring non-target-site resistance remain largely unknown (Suzukawa et al., 2021). Both types of herbicide resistance mechanisms can be found in a single weed population and even within a single individual (Ghanizadeh et al., 2022).

*Lolium multiflorum* is a winter annual, outcrossing, diploid (2n = 14) plant species native to the Mediterranean basin (Humphreys et al., 2009). It can drastically reduce crop yield if left unmanaged. In the winter cereal wheat, for example, competition with *L. multiflorum* can cause yield losses in the order of 50% (Appleby et al., 1976). Furthermore, when *L. multiflorum* grows where crops grown for seed are cultivated, it can contaminate commercial seed lots, resulting in long-distance spread of weed seeds. This uncontrolled movement of *L. multiflorum* genotypes can result in admixture with natural grassland populations of interfertile plant species (e.g., *L. perenne*), impacting the genetic diversity across the landscape (Meade et al., 2020).

Since 1987, many populations of *L. multiflorum* have evolved herbicide resistance (Heap, 2022). Resistance to herbicides that inhibit acetyl-CoA carboxylase (ACCase) has been notably frequent around the world, with 35 populations from 10 countries reported to date, in areas where repetitive use of ACCase inhibitors occur. ACCase inhibitors selectively control grass species, allowing their direct application to broadleaf crops without considerable damage to the crop. Furthermore, because wheat can metabolize some ACCase inhibitors to less active compounds (Yu & Powles, 2014), this class of herbicide has been widely used in this major staple crop. These characteristics, in addition to their safe environmental and human safety (EPA, 2020), has led to the broad adoption of ACCase inhibitors.

ACCase is an important enzyme in plant metabolism, being responsible for the formation of malonyl-CoA from the carboxylation of acetyl-CoA. Depletion of malonyl-CoA, which is the substrate for fatty acid biosynthesis in the plastids, results in loss of cell membrane integrity because of the lack of lipids and other secondary metabolites, ultimately resulting in cellular leakage and plant death (reviewed by Devine, 2002). ACCase inhibitors target the carboxyl transferase (CT) domain of the enzyme (Yu et al., 2010). The specific molecular mechanisms conferring resistance in *L. multiflorum* populations are often unknown. However, in general, ACCase resistance is manifested either by amino acid substitutions in key residues of the CT domain, or enhanced herbicide metabolism, whereas the former seems to be the most frequently reported molecular mechanism (Powles & Yu, 2010). Several non-synonymous single nucleotide polymorphisms (SNPs) have been identified in weed populations conferring resistance to ACCase inhibitors (Murphy & Tranel, 2019). The resultant amino acid substitutions are located at or near the active site of ACCase, altering the molecular interactions with herbicides. For example, when an isoleucine is substituted by a leucine at position 1781, the anchoring of the methyl or ethyl group (depending on herbicide chemical class) of the herbicides is altered. Another example is when amino acid substitutions are observed at position 2078 of the ACCase, alterations in nearby side chains of amino acids change the conformation necessary for binding (Yu et al., 2010).

In this work, we examined 14 *L. multiflorum* populations from various cropping systems in northwest Oregon, USA. The objective was to elucidate the mechanisms of herbicide resistance in populations from agricultural fields exposed to strong human-driven selection pressure with herbicides, and to examine the repeatability of herbicide resistance evolution in this important weed species. We investigated the ACCase resistance patterns and used long-read DNA sequencing to elucidate their genetic relationships. Our findings highlight the astonishing genetic diversity of *L. multiflorum* populations and the repeatability of ACCase inhibitor resistance evolution in agricultural fields, and we provide a discussion of basic and practical implications of our results.

## MATERIALS AND METHODS

### Plant material

Field populations of *L. multiflorum* were collected from wheat and seed crop fields in 2017-2018 (Table S1) as part of a broader herbicide resistance survey (Bobadilla et al., 2021). Weed management in this cropping system is performed primarily with herbicides from various mechanisms of action. Crop rotations can also be part of an integrated weed management strategy, with a grass crop (e.g., annual or perennial ryegrass grown for seed or wheat) typically following a broadleaf crop (e.g., clover grown for seed, radish, or meadowfoam). Briefly, seed from 25-30 mature plants were individually sampled and later pooled in approximately equal quantities. Detailed information of sampling procedures is available elsewhere (Bobadilla et al., 2021).

A total of 14 *L. multiflorum* field populations were selected for this research (Figure 1) based on an initial screening with a commercial dose of ACCase inhibitors. In addition to the field populations, we also included a cultivated variety of *L. multiflorum* (popGulf) that we expected to be susceptible to all herbicides, as well as a previously characterized ACCase inhibitor resistant population, popPRHC (Brunharo & Hanson, 2018). To minimize maternal effects from field populations, we generated a bulked population (B_1_) to be used for resistance level quantification. The B_1_ population was produced as follows. Approximately 30 seeds from each population were germinated, transplanted to pots filled with commercial potting media, and grown to maturity in a greenhouse. Individuals were allowed to cross pollinate within each population. Cross pollination between populations was avoided by both isolating populations in separate greenhouses and temporal isolation during pollen dispersal.

**Figure 1.**
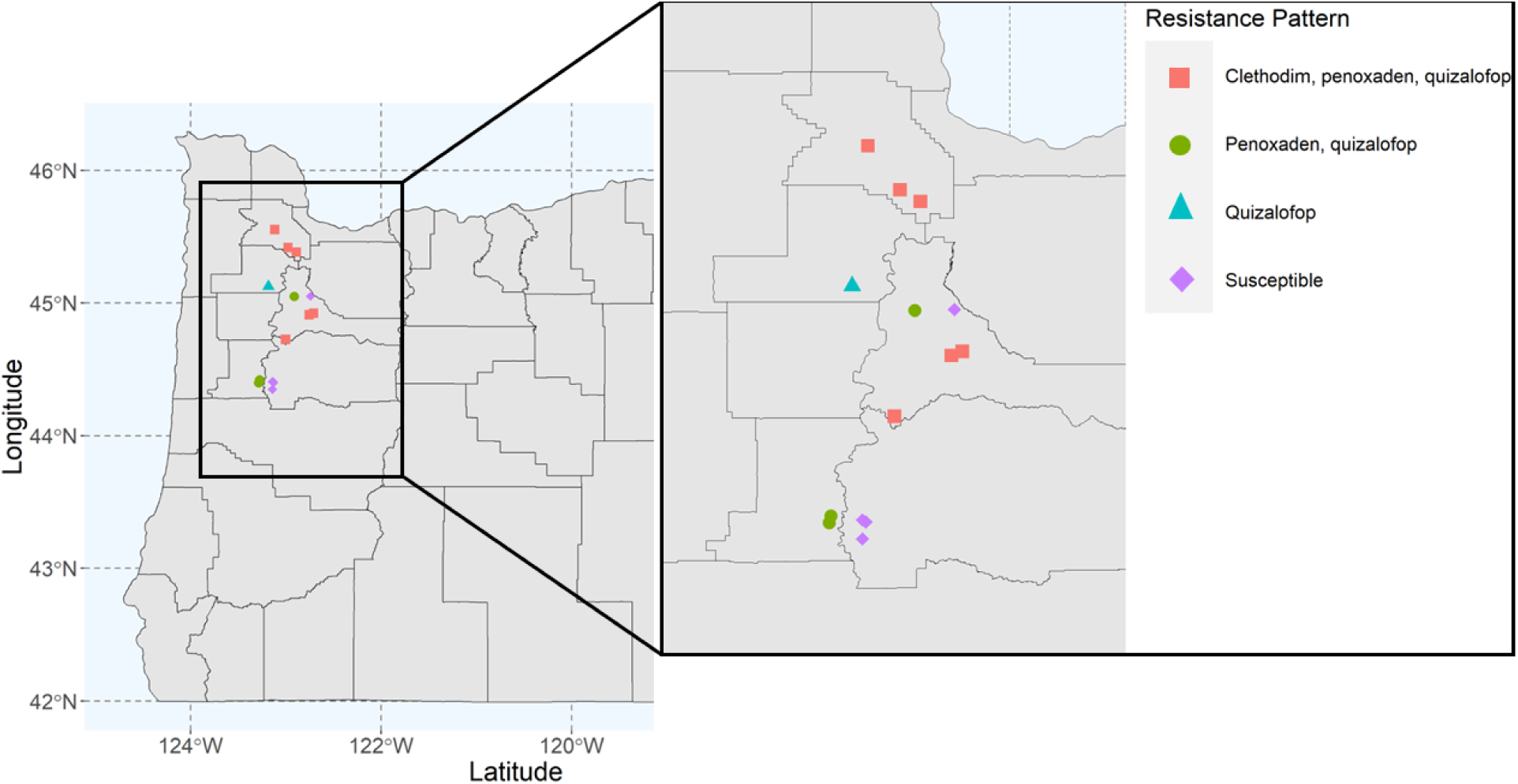
Locations in Oregon, USA where *Lolium multiflorum* populations were collected. Blue squares represent susceptible populations, yellow circles represent resistance to one ACCase inhibitor, green triangles represent resistance to two ACCase inhibitors, and red diamonds represent resistance to three ACCase inhibitors.

### Dose-response experiments

Dose-response experiments were conducted with the B_1_ populations to identify and quantify resistance to ACCase inhibitors. Clethodim, pinoxaden, and quizalofop-P-ethyl (quizalofop henceforth) were chosen to represent herbicides from the three distinct ACCase inhibitor chemical groups (cyclohexanediones, phenylpyrazolin, and aryloxyphenoxy-propionates, respectively). These herbicides were selected given their importance in the cropping systems where *L. multiflorum* populations were collected. Clethodim is an herbicide commonly used in broad leaf seed crops for grass weed control, typically applied postemergence multiple times during the growing season. Pinoxaden is a selective, postemergence grass herbicide for wheat that can be used at different growth stages throughout the growing season. Finally, quizalofop was selected because of the predicted increased importance of this herbicide in wheat, because crop varieties resistant to this active ingredient are being released in multiple locations across the United States (Hildebrandt et al., 2022). Other herbicides from the same chemical group of quizalofop have been extensively used in the region for *L. multiflorum* control, particularly diclofop-methyl. Therefore, quizalofop is expected to reflect resistance to herbicides in the aryloxyphenoxy-propionates chemical group.

Plants from the B_1_ populations were germinated in Petri dishes in a growth chamber programmed for 14/10h (day/light) and approximately 200 µmol m^-2^ s^-1^ of photosynthetically active radiation (PAR). Seedlings were transplanted to pots, one per pot, filled with a commercial potting media and transferred to a greenhouse (14/10h) for the remainder of the studies. When plants reached the BBCH-22 growth stage (two side tillers visible, approximately 10 cm in height), dose-response experiments were conducted. Dose-response studies are designed to expose plants to increasing rates of herbicides to estimate the dose at which growth is reduced by 50% (GR_50_). Estimates are generated for the suspected resistant as well as a known susceptible, then compared to estimate the resistance index. Our dose-response studies had eight doses: zero (nontreated control), 0.125X, 0.25X, 0.5X, 1X, 2X, 4X, 8X, where X represents the recommended field rate, in four replications. The field rates for clethodim, pinoxaden, and quizalofop were 102, 60.3, and 92.5 g of active ingredient (a.i.) ha^-1^, respectively. Visual injury data, ranging from 0-100 (where 0 represents absence of injury and 100 represents complete control), were collected 28 days after treatments, followed by sampling of aboveground plant biomass by cutting plants at the soil surface and drying in a forced-air oven. The dose-response experiments were repeated. In total, we generated 96 dose-responses (three herbicides × 16 populations × two experimental runs).

### Amplicon-seq preparation and sequencing

Resistance to ACCase inhibitors has been previously reported to be conferred by amino acid substitutions in the CT domain of the target site. Therefore, we hypothesized that ACCase inhibitor resistance in *L. multiflorum* populations was caused by single nucleotide polymorphisms in the *ACCase* gene. To test this hypothesis, we sequenced the entire CT domain of *ACCase* from 97 resistant and susceptible *L. multiflorum* individuals (Table S2) to elucidate the role of polymorphisms in the resistance patterns, as well as the genetic relationship among sampled individuals and populations.

As the first step, populations were phenotyped for resistance to clethodim, pinoxaden, or quizalofop before sequencing. Seeds (collected from the field) from each population were germinated and grown as previously described. Plants were grown to approximately six to ten tillers, and tillers were split into four clones. Single tillers were transplanted to individual pots and allowed to recover in a greenhouse. This step was taken to test a single genotype with all three ACCase inhibitors of interest (i.e., clethodim, pinoxaden, and quizalofop), in addition to an untreated control, allowing us to assess the resistance patterns at the individual level. Plants were treated with twice the recommended field rate when they reached the BBCH-22 (Hess et al., 1997) growth stage. Our goal was to sequence 10 individuals per population. However, because some tillers did not survive transplant, phenotyping with all three ACCase inhibitors was not possible. Ultimately, we sampled 4-10 plants per population (Table S2). Leaf tissue was collected from each individual genotype the day prior to herbicide treatment, and kept in a -80 C freezer until analysis. Approximately four weeks after treatment, plants were phenotyped as alive or dead (0 = dead, 1 = alive). DNA was extracted from *L. multiflorum* samples using a commercial DNA extraction kit (Wizard HMW DNA Extraction Kit, Promega Corporation, Madison, WI, USA).

We developed primers to amplify the entire coding sequence of the CT domain of ACCase. We focused on this region of the *ACCase* because the CT domain is where all ACCase inhibitors bind (Zagnito et al., 2001). In addition, known amino acid substitutions conferring resistance to this herbicide mechanism of action are in the CT domain. To design primers, we aligned the DNA sequence of the *ACCase* CT domain from *E. crus-galli* (Xia et al., 2016) to the *L. multiflorum ACCase* sequence from White et al. (2005) (NCBI accession number AY710293.1) using Geneious v11.0.14.1 (www.geneious.com). This step was performed to identify the CT domain coding sequence in *L. multiflorum*. Primers were designed with the Primer3 (v.2.3.7) feature of Geneious (Untergasser et al., 2012). Several primer pairs were tested with *L. multiflorum* DNA and under different PCR conditions, and chosen based on results of electrophoresis analysis and the expected number of fragments and size. The primer pair utilized for the amplicon-sequencing experiment was 5’-AGGGAGCACTGTTGTGGATG-3’ (forward) and 5’-GTTCTCCCTCCAGGCAACAA-3’ (reverse), amplifying a 2,411 bp region of *ACCase* containing the CT domain (size based on reference *ACCase* sequence from White et al., 2005).

Amplicon generation and library preparation were performed at the Functional Genomics Facility, Biotechnology Center, at the University of Illinois Urbana-Champaign using a commercial kit following the manufacturer’s recommendations (Part number 101-599-700, Pacific Biosciences, Menlo Park, CA, USA). Briefly, genomic DNA sample concentration was normalized to 0.175 ng µL^-1^ with a Qubit (Invitrogen, Waltham, MA, USA). Amplicons were amplified for each sample by PCR with forward and reverse uniquely barcoded primers as follows: initial denaturation for 3 min at 95 C, then 27 cycles of denaturation for 30 s at 98 C, annealing for 30 s at 60 C, and extension for 5 min at 72 C. Reactions were visualized in an agarose gel. Finally, amplicons were pooled in equal amounts and purified with magnetic beads (AMPure XP, Brea, CA, USA).

PacBio long-read sequencing library preparation and sequencing was performed at the High-Throughput Sequencing and Genotyping Facility, Biotechnology Center, at University of Illinois Urbana-Champaign. The pooled amplicons were converted into a PacBio library with the SMRTbell Express Template Prep kit version 2.0 (Pacific Biosciences, Menlo Park, CA, USA). The library was quantitated with Qubit and run on a Fragment Analyzer (Agilent, Santa Clara, CA, USA) to confirm the presence of DNA fragments of the expected size. The library was sequenced on one SMRT cell 8M on a PacBio Sequel IIe with 30 h movie time. The circular consensus analysis was in real time in the instrument with SMRT Link V10.1 using 99.9% accuracy (HiFi reads).

### Data analysis

Visual injury and biomass data were analyzed with log-logistic regressions. First, data across experimental runs were pooled after passing the Levene’s homogeneity test (LeveneTest function from the *car* package in R) (Fox & Weisberg, 2019). Dose-response experiments were fit into three parameter log-logistic regressions (*fct = LL.3*) using the *drc* package in R (Ritz et al., 2015). For each population, the GR_50_ (herbicide dose required to reduce plant growth by 50%) was estimated and compared to the population Gulf (popGulf, known susceptible). The resistance index (RI; the ratio between the GR_50_ from a resistant population compared to the known susceptible) was calculated to estimate the magnitude of the resistance phenotype. The *predict* function in R was used to generate confidence intervals around the mean values generated by the log-logistic models. Finally, data were plotted with ggplot2 (Wickham, 2016). Populations were classified as resistant if the RI was greater than 2.

The HiFi amplicon data were analyzed with PacBio tools (https://github.com/PacificBiosciences/pbbioconda). Data were demultiplexed and sequencing primers and barcodes removed with the *lima* algorithm. The *pbaa* pipeline was followed to analyze the dataset. Data from each sample was clustered to generate consensus sequences using the *L. multiflorum ACCase* coding sequence as the guide reference and default filters were applied. Fasta files containing clustered sequences that passed the default filtering step were split to one sequence per file to represent each haplotype. After conversion from fasta to fastq using a Perl script (https://github.com/ekg/fasta-to-fastq/blob/master/fasta_to_fastq.pl), sequences were aligned to the reference ACCase sequence, converted to *bam*, and sorted using samtools (Li et al., 2009).

The HaplotypeCaller pipeline (Van der Auwera & O’Connor, 2020) was used to generate a SNP and insertion/deletion dataset for the analyses that follow. The *HaplotypeCaller* algorithm was used with each *bam* file, using the *ACCase* as reference, and the parameters --*min-pruning 0 and -ERC GVCF* were used to generate an individual variant dataset for each sample. Second, we used *CombineGVCFs* to combine the individual variant file from each sample into a single cohort VCF file. Finally, a joint genotyping step (*GenotypeGVCFs*) was performed with the combined VCF file, outputting a final VCF that accounts for population-wide information to improve genotyping sensitivity.

A principal component analysis (PCA) was performed to obtain an initial examination of genetic variability in the haplotypes both within and between *L. multiflorum* populations. Because PCA does not require any assumption of the underlying population genetic model, this method was used to infer the relationships between individuals based on our *ACCase* variant dataset (Jombart et al., 2010). The R package SNPRelate (Zheng et al., 2012) was used to import the VCF file, generate eigenvectors, and calculate the variance proportion for each principal component, followed by plotting with ggplot2 (Wickham, 2009). Variant information was manually inspected in Geneious to identify polymorphisms in the *ACCase* gene known to confer resistance to ACCase inhibitors (Murphy & Tranel, 2019).

To further dissect the genetic relationships among *L. multiflorum* individuals and understand the evolutionary origin of the resistance alleles, we generated a maximum-likelihood phylogenetic tree with the *ACCase* haplotype sequence. Multiple sequence alignment was performed with MAFFT (v.7.487) (Katoh et al., 2002) (*--reorder --maxiterate 1000 --retree 1 --genafpair*), and the resulting alignment output was subjected to RAxML genetic tree inference algorithm with Felsenstein bootstrapping with 200 replicates (*--all --model LG+G8+F --tree pars {10} --bs-trees 200*) (Kozlov et al., 2019). A maximum-likelihood tree was generated with the R package *ggtree* (Yu et al., 2017). To facilitate visualization of divergence among haplotypes from the different populations and infer the involvement of gene flow in the spread of herbicide resistance alleles, we constructed a phylogenetic tree and labeled the tip based on the population name. The role of isolation by distance in the evolution of herbicide resistance was assessed with a Mantel Test implemented in the r package *adegenet* (Jombart, 2008).

We investigated the diversity of haplotypes within each population by calculating the expected heterozygosity, nucleotide diversity, and number of polymorphisms using the populations module from *Stacks* (Rochette et al., 2019). The multiple alignment file generated with MAFFT was used as input to DnaSP (v6) (Rozas et al., 2017) to calculate molecular population genetic statistics such as average length of the sequences, number of haplotypes, and haplotype diversity (Nei, 1987).

During data analysis, we observed that several resistant individuals did not exhibit any known resistance-endowing polymorphisms that would lead to amino acid substitutions in the *ACCase*. However, we identified several non-synonymous SNPs, as well as long indels, in the coding sequence. We therefore conducted an association study to test the hypothesis that novel mutations were responsible for the resistance phenotypes. First, as a proof of concept, we selected all haplotypes that exhibited SNPs known to confer ACCase resistance and performed an association study with the Blink model (Huang et al., 2018) implemented in GAPIT (Lipka et al., 2012). The VCF file obtained from previous analysis was used in this study after conversion to a HAPMAP format (Gibbs et al., 2003) with TASSEL (Bradbury et al., 2007). After confirmation of the reliability of this approach, we performed an association analysis with amplicons containing non-synonymous SNPs not previously reported to confer resistance to ACCase inhibitors.

## RESULTS

Dose-response experiments with clethodim, pinoxaden, and quizalofop revealed widespread ACCase inhibitor resistance in *L. multiflorum* populations from sampled fields. Three and six of the field populations, respectively, exhibited cross-resistance to two and three of the herbicides analyzed (Table S1). Cross-resistance is defined as resistance to herbicides from more than one chemical group within a mechanism of action. Surprisingly, only one population exhibited resistance to a single herbicide chemistry. Four of the field populations exhibited susceptibility to all herbicides tested. As expected, the known susceptible cultivar of *L. multiflorum* (popGulf) was susceptible to all herbicides, and the known resistant population (popPRHC) survived penoxaden and quizalofop. The RIs varied considerably depending on the herbicide (Table S3). Resistant populations exhibited RIs of 2-13 to clethodim, indicating that they require up to 13 times more clethodim than the reference susceptible to cause 50% of biomass reduction. This estimate was 5-20 for pinoxaden, and 2-37 for quizalofop. We also observed large variation in response to the herbicide within doses, suggesting the populations are segregating for multiple genotypes that may or may not contain *ACCase* resistance alleles.

The amplicon library generated nearly 2M PacBio HiFi reads, with mean amplicon length of 2,762 bp. Our SNP discovery pipeline resulted in the resolution of 202 haplotypes from 97 *L. multiflorum* individuals. The larger-than-expected number of haplotypes found can be attributed to the discovery of tetraploid individuals in our collection. Our SNP discovery pipeline resulted in a VCF file with 201 variants, of which 12 were insertion/deletions (insertions were 1-24 bp-long; deletions were 1-31). We observed that 116 of the SNPs were transitions (i.e., A to G, or C to T).

We were particularly interested in surveying SNPs occurring at coding positions 1781, 1999, 2027, 2041, 2078, 2088, and 2096 of the ACCase enzyme, because these positions have been previously associated with resistance to ACCase inhibitors. Of the SNPs identified, eight resulted in amino acid substitutions at position 1781, 18 at position 2027, 28 at position 2041, and 16 at position 2078. The population that exhibited the greatest diversity in resistance-endowing polymorphisms was population pop19, where all individuals were ACCase inhibitor-resistant and had amino acids substitutions at positions 1781, 2027, 2041, and 2078 (Table S2). Not surprisingly, these amino acid substitutions did not occur in the same haplotype. In fact, no haplotype with more than one resistance-endowing SNP was identified from our dataset. Population pop92, similarly, had a high proportion (7 out of 8) of individuals exhibiting resistance, with polymorphisms at positions 2027, 2041, and 2078. Out of the 97 individuals sequenced and phenotyped, 57 exhibited resistance to at least one ACCase inhibitor (Table S2). A few individuals that survived a 2 × application of ACCase inhibitor did not exhibit any known amino acid substitution (a total of 17; Table S2). In addition, several individuals exhibited resistance patterns that have been previously established, whereas some others did not (Table S4). In total, 70 of the 202 haplotypes had resistance-endowing SNPs. These results indicate that individuals heterozygous for the ACCase allele can survive to lethal herbicide rates, corroborating with multiple reports from the literature (Yu et al., 2013).

Molecular population genetic summary statistics suggest a large diversity of haplotypes among *L. multiflorum* individuals and populations (Table 1). The length of the *ACCase* sequence mapped varied from 2,629 to 2,665 bp, indicating the existence of haplotypes with multiple indel events. DnaSP analysis indicated the total number of unique haplotypes identified in the dataset totaled 83, which is also reflected by the high haplotype diversity value (H_d_) of 0.989. Note that in Table 1 the total number of haplotypes does not equate to the sum of the haplotypes from all populations, because some of the haplotypes were shared among populations. Finally, the nucleotide diversity (π) estimate varied from 0.070 to 0.152, indicating a large number of polymorphism events. The expected heterozygosity data was displayed in a violin plot along with the population average (Figure S1). Expected heterozygosity varied from as low as 0.065 to as high as 0.153, where larger values suggest more *ACCase* diversity. Given an herbicide application can drastically reduce the population size, we performed a Pearson correlation analysis between the expected heterozygosity and the number of herbicide groups to which the population was resistant. We expected that a population resistant to a greater number of herbicides would have less diversity because of the repeated bottleneck events from herbicide applications and consequent reduction of diversity. However, we observed no correlation between expected heterozygosity and herbicide resistance (P = 0.62).

**Table 1.**
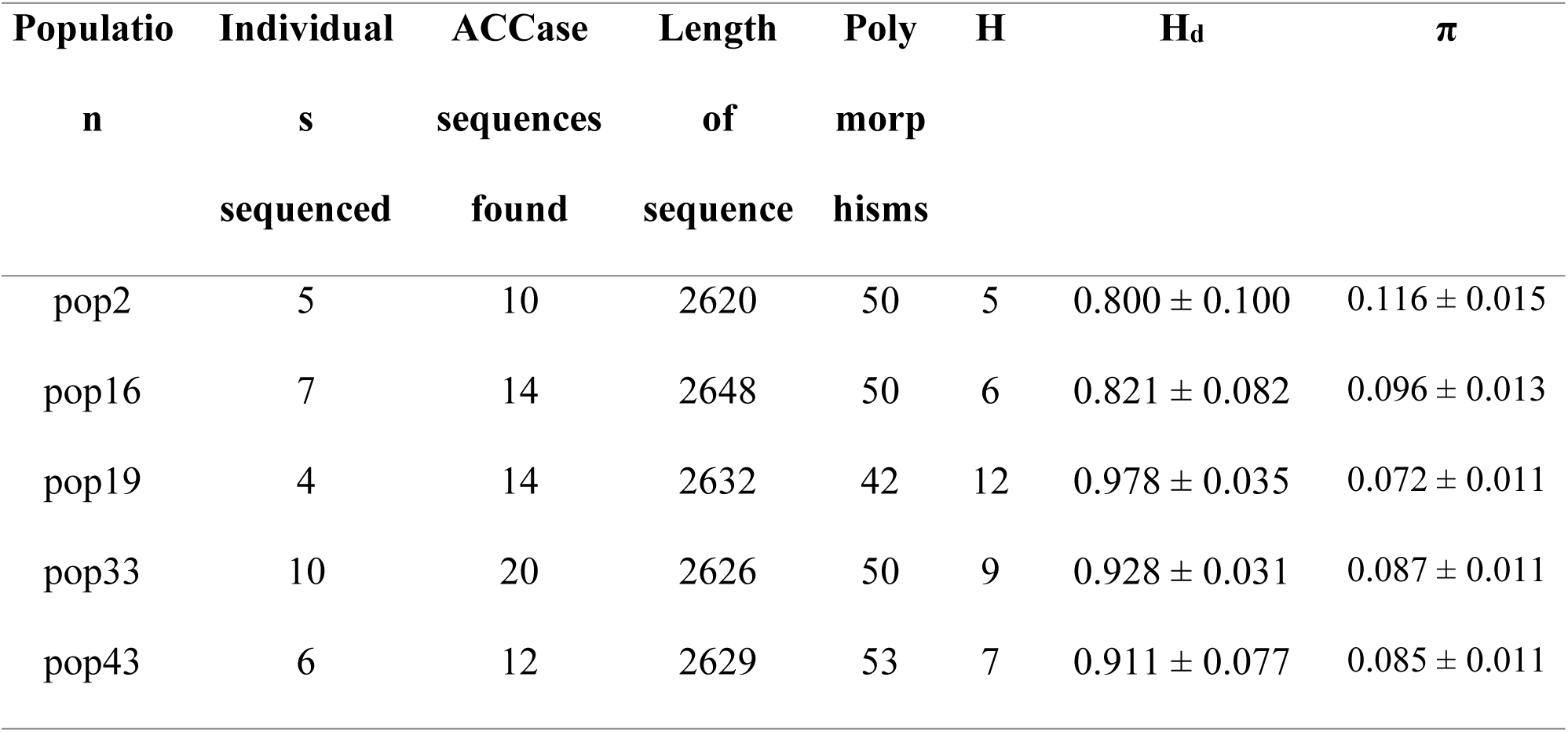
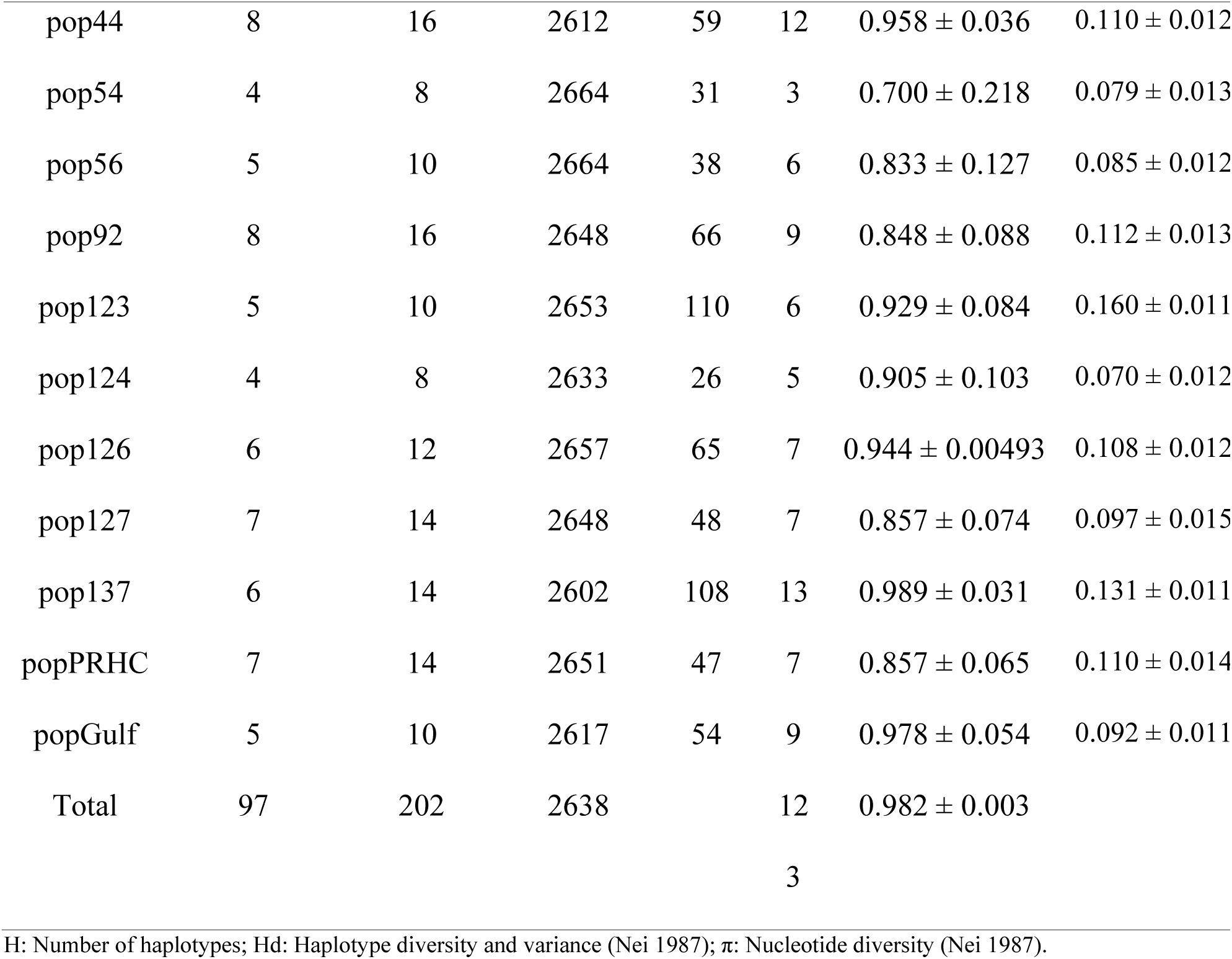
Summary of population genetics estimates.

Principal component analysis of all populations revealed large genetic variation among the *L. multiflorum* individuals analyzed. When all populations were included in the analysis, PC1 and PC2 explained, respectively, 30 and 11% of the variation (Figure 2, Upper Panel). Individuals containing known resistance-endowing polymorphisms were separated primarily by PC2 (Figure 2, Lower Panel). We did not observe any patterns in how individuals with known ACCase resistance SNPs clustered in the PCA.

**Figure 2.**
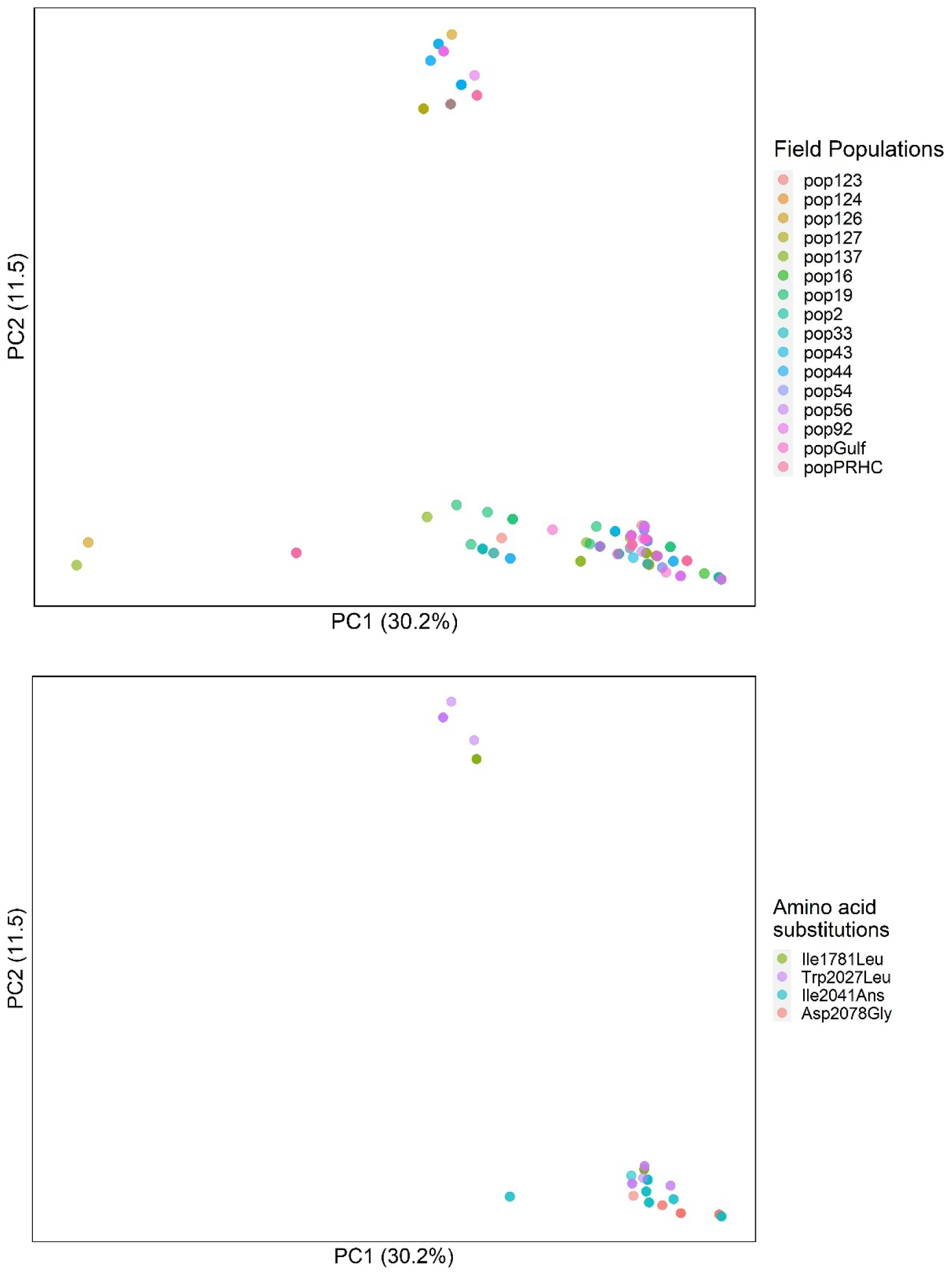
Principal component analysis of *Lolium multiflorum* populations susceptible and resistant to ACCase inhibitors. Upper panel: PCA with all individuals. Lower panel: PCA with resistant individuals only (67 haplotypes).

The maximum-likelihood tree generated corroborated with the population genetics statistics that multiple haplotypes exist in our analysis. Figure 3 exhibits each individual haplotype, labeled based on whether they contain resistance-endowing SNPs. This analysis indicated there is a strong divergence in the genetic relationships among the haplotypes, supported by bootstrapping replicates. We created a similar figure with the same underlying dataset, however with the tips of the phylogenetic tree labeled according to the population from which the haplotype originated (Figure S2). We observed widespread admixture of *ACCase* among the populations analyzed. These conclusions are supported by the Mantel test, where we did not observe that genetic variation was associated with spatial isolation of populations (Figure S3).

**Figure 3.**
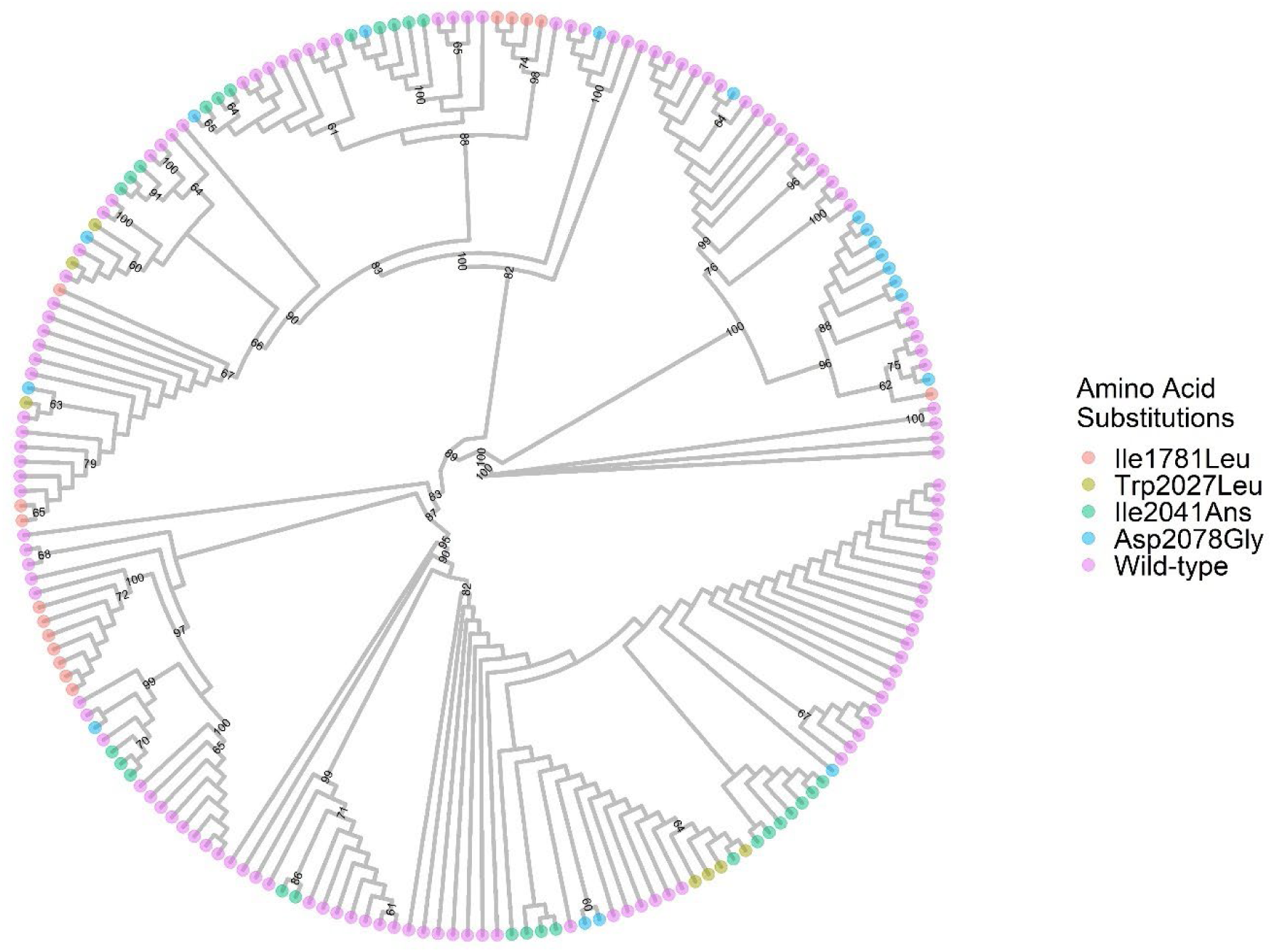
Best-scoring maximum likelihood phylogenetic tree of *Lolium multiflorum* based on *ACCase* sequence. Numbers indicate bootstrapping values (values smaller than 60 were omitted). Tips were colored based on whether the haplotype had an amino acid substitution at positions 1781, 2027, 2041, or 2078, or was wild type at these positions.

Because 17 individuals phenotyped were ACCase inhibitor resistant but did not exhibit any resistance-endowing amino acid substitution, we performed an association study to identify if other polymorphisms in the gene could be involved in the resistance phenotype. As a proof of concept, first we only included haplotypes containing known amino acid substitutions, and regressed them with a binary response variable as dead or alive. The results indicated that the statistically significant SNPs were the non-synonymous mutations at positions 1781, 2027, 2041, and 2078 (Figure S4, Upper Panel). Next, we performed an association analysis including only resistant individuals that did not exhibit any resistance-endowing SNPs. This association analysis indicated that unknown polymorphisms found in the *ACCase* were not associated with the resistance phenotype (Figure S4, Lower Panel).

## DISCUSSION

Our research aimed to resolve the complexity in ACCase inhibitor resistance patterns in *L. multiflorum*, and to elucidate the evolutionary relationships among populations with distinct responses to this group of herbicides. We observed a remarkable genetic diversity in *ACCase*, and our data suggest multiple origins of *ACCase* resistance alleles despite the relatively small geographic region from which the populations were collected. In addition to finding resistance-endowing polymorphisms in many different *ACCase* haplotypes, we also observed widespread admixture of resistance alleles among the different populations. This large genetic diversity observed in the coding sequence of *ACCase* plays a crucial role in the repeated evolution of herbicide resistance in *L. multiflorum*. Although we only focused on the CT domain of ACCase, other studies have also concluded that *L. multiflorum* exhibits large genetic diversity in other genomic regions (Karn & Jasieniuk, 2017; Tamura et al., 2022).

Given the large number of distinct haplotypes identified in the field populations, it can be inferred that, at the nucleotide sequence level, both convergent and parallel evolution have played an important role in the selection of resistant populations of *L. multiflorum*. It is unclear, however, the evolutionary mechanisms by which herbicide resistance became widespread in the region. Evolution of herbicide resistance can occur *via* the rise of new beneficial mutations, selection of beneficial mutations from standing genetic variation, or immigration *via* gene flow (Lee & Coop, 2017). Because we only sequenced the coding region of the CT domain of ACCase, we were unable to establish the underlying evolutionary mechanism. It is important to note that the populations sampled are from agricultural systems, where human-mediated gene flow is constant and intense. Movement of farm equipment and trade of seed lots contaminated with weed seeds, for example, could facilitate the admixture among populations from many parts of the region, country, and globe. Therefore, it is possible that each resistance allele, before introduction to Oregon, has experienced distinct histories of selection pressure and evolved via different evolutionary mechanisms. Regardless of the evolutionary mode of adaptation to ACCase inhibitors, our results support the hypothesis that resistance evolution to ACCase inhibitors in *L. multiflorum* happened multiple times because we found the same resistance-endowing SNPs on different haplotypes, and widespread admixture is supported by PCA, phylogenetic, and isolation-by-distance analyses. These results are expected of an obligate-outcrossing weed species with large standing genetic variation. The fact that we did not observe correlation between expected heterozygosity and the number of herbicides the population is resistant to may indicate that recurrent gene flow could minimize the effect of bottleneck events, replenishing the genetic diversity lost after an herbicide application.

It has been previously suggested that the resistance patterns observed in grasses could be explained by amino acid substitutions in ACCase (Powles & Yu, 2010). For instance, it is generally assumed that amino acid changes at position 1781 will confer resistance to herbicides in all three chemical groups of ACCase inhibitors based on previous work in *Alopecurus myosuroides* (Petit et al., 2010) and *Lolium rigidum* (Zhang & Powles, 2006), while substitution at position 2078 confers resistance to all chemical groups in *A. myosuroides* (Petit et al., 2010), *L. multiflorum* (Kaundun, 2010), *L. rigidum* (Yu et al., 2007), and *P. paradoxa* (Hochberg et al., 2009). Our research indicates that target-site alteration alone is not sufficient to accurately predict the resistance phenotype in *L. multiflorum*. For instance, we identified 15 individuals exhibiting an amino acid substitution at position 2041 (Table S4), where one was resistant to only quizalofop, two to clethodim and quizalofop, four to pinoxaden and quizalofop-ethyl, and eight to clethodim, pinoxaden, and quizalofop. Conversely, our results corroborate with previous data that an amino acid substitution at position 2078 confers resistance to all chemical classes, based on observation from 10 individuals with this genotype (Table S4). The complexity in resistance patterns could be explained by additional polymorphisms in the ACCase gene (other than those known to confer ACCase inhibitor resistance). These additional polymorphisms could alter the structure of the binding pocket, reestablishing the inhibition of the enzyme by herbicide of a specific chemical class. This compensatory mechanism has been suggested elsewhere (Yu et al., 2010). Unfortunately, we were not able to identify these additional polymorphisms in our association analysis (Figure S4, Upper Panel), likely because the number of individuals with unusual resistance patterns was small. Alternatively, co-existence of non-target mechanisms could confound cross-resistance patterns.

In addition to being a weed in agricultural systems, *L. multiflorum* is also a plant species used as cover crop and pasture (OECD, 2022). Breeding programs have released tetraploid varieties with beneficial agronomic traits. Our sequencing analysis identified four tetraploid individuals. Out of the 16 haplotypes, four had known ACCase resistance-conferring SNPs. The origin of these polymorphisms is unclear. The phylogenetic analysis suggests that these haplotypes have close evolutionary origins with other non-resistant, diploid individuals, suggesting that the resistance alleles might have been present in the breeding programs prior to chromosome duplication and release of the varieties. Because tetraploid cultivars are not compatible with their diploid counterparts (Schmitz et al., 2020), if evolution happened after release of cultivated varieties, we would expect to see greater differentiation and separation from other haplotypes, which is not the case.

When new introductions occur in natural areas, it is expected that the genetic diversity will reduce due to bottlenecks, and with time clear differentiation can be observed compared to the source population (Amsellem et al., 2001). Remarkably, our data suggests that, despite evidence of gene flow among populations, the diversity of resistance haplotypes is maintained in *L. multiflorum*. This could be explained by two aspects. First, the small geographic region associated with a dynamic cropping system facilitates recurrent introductions of weedy *L. multiflorum* along with cultivate *L. multiflorum via* seed contamination. Secondly, it is possible that a limited number of seed from herbicide resistant individuals is introduced to a new area (creating a bottleneck); however, given weedy *L. multiflorum* may hybridize to cultivated *L. multiflorum*, the genetic bottleneck is promptly minimized and resistance (and other weedy) genes are quickly introgressed to local populations (Matzrafi et al., 2021). More research is needed to elucidate these dynamics in *L. multiflorum*.

Weed populations have been shown to exhibit large genetic variability, diversity, and structure at the landscape level. Comont et al. (2020) have shown that many populations of *A. myosuroides* from the United Kingdom exhibited TSR to ACCase inhibitors conferred by different mutations, with frequencies varying from <0.1 to >40%. Similarly, Kersten et al. (2023) evaluated the ACCase alleles from 27 *A. myosuroides* populations from across Europe, and observed that most populations had at least two distinct TSR mutations. Other authors have observed large degree of admixture in herbicide resistant populations from agricultural fields at the landscape level (Kuester et al., 2015; Okada et al., 2013; Dixon et al., 2020; Ravet et al., 2020). Conversely, there are examples where herbicide resistant weed populations have admixture in some populations, but not others (Lawrence et al., 2017; Kreiner et al., 2019; Küpper et al., 2018). The genetic characteristic of a plant population is an interplay of genetic drift, gene flow, and natural selection (Eckert et al., 2008). These processes will dictate the level of genetic diversity observed. In agricultural systems, not only species biology, environment, and life history, but also weed management practices locally and at the landscape level will influence the resulting genetic makeup of populations.

The fact that we identified individuals that exhibited ACCase inhibitor resistance, but did not have polymorphisms in the target site, suggests that non-target site resistance mechanisms are also involved. Non-target site resistance to ACCase inhibitors in *L. multiflorum* has been documented multiple times. For instance, Cocker et al. (2001) found that resistant populations exhibited an enhanced ability to metabolize the herbicides, likely mediated by glutathione S-transferases. Han et al. (2014) found that populations of *L. rigidum*, a close relative of *L. multiflorum*, exhibited mixed resistance mechanisms to ACCase inhibitors, where the amino acid substitution at position 2041 was predominant in their collection.

Timely herbicide resistance detection is crucial to allow farmers to adjust management practices to the presence/absence of herbicide-resistant weed populations, as well as to minimize the spread of resistant populations. There have been efforts to identify SNPs conferring resistance or linked to herbicide-resistance alleles, with the objective of developing genetic markers for quick herbicide resistance detection, such as KASP assays (Mendes et al., 2020). Given its genetic variability and multiple nucleotide substitutions that can confer herbicide resistance, multiple detection assays would have to be developed and run for each sample, increasing costs and time. Kersten et al. (2022) developed a pool-seq workflow to identify target site resistance in *Alopecurus myosoroides* populations from field populations in Germany to assist with weed management decisions. The pool-seq approach is more likely to be feasible for capturing multiple herbicide resistance-endowing polymorphisms compared to KASP assays. A pitfall for using detection methods based on DNA sequences is that the variants conferring the phenotype must be known. This will be particularly challenging for weed populations with mixed resistance mechanisms, because the genetic bases for non-target site resistance are rarely known (Suzukawa et al., 2020).

There are many questions that remain unanswered. Our haplotype-based analysis identified DNA sequences with several polymorphisms, and large insertions and deletions were observed. It remains to be seen the effects of these alterations on enzyme kinetics with the natural substrate and herbicides. These mutations were observed in the heterozygous and homozygous state (data not shown), suggesting the enzyme remained functional. However, it is unclear to what extent they could result in ecological penalty under certain growing conditions. Our association analysis was not powerful enough to detect the involvement of new polymorphisms in *ACCase* that could confer herbicide resistance. However, this hypothesis should be further investigated with a larger panel of plants and controlled crosses. It also remains unclear what are the non-target site resistance mechanisms involved in the studied populations; therefore, future research could address the relative contribution of each resistance mechanisms.

In summary, our results identified large *ACCase* variability in *L. multiflorum* populations collected from agricultural fields. Cross-resistance to ACCase inhibitors was widespread, and the resistance patterns across the different chemical groups seem complex and likely governed by multiple herbicide-resistance mechanisms. Known herbicide-resistance polymorphisms in the *ACCase* gene likely evolved multiple times in the populations studied.

## DATA ACCESSIBILITY

Data for this study are available under BioProject ID PRJNA993136.

## Acknowledgements

We would like to thank the staff at the Functional Genomics Unit and the Roy J. Carver Biotechnology Center at the University of Illinois Urbana-Champaign for their technical assistance in the generation of amplicon sequencing data. Funding for this project was provided by the College of Agricultural Sciences at The Pennsylvania State University and Oregon State University.

