## Supplemental file 1 for "Repeated evolution of herbicide resistance in *Lolium multiflorum* revealed by haplotype-resolved analysis of acetyl-CoA carboxylase"

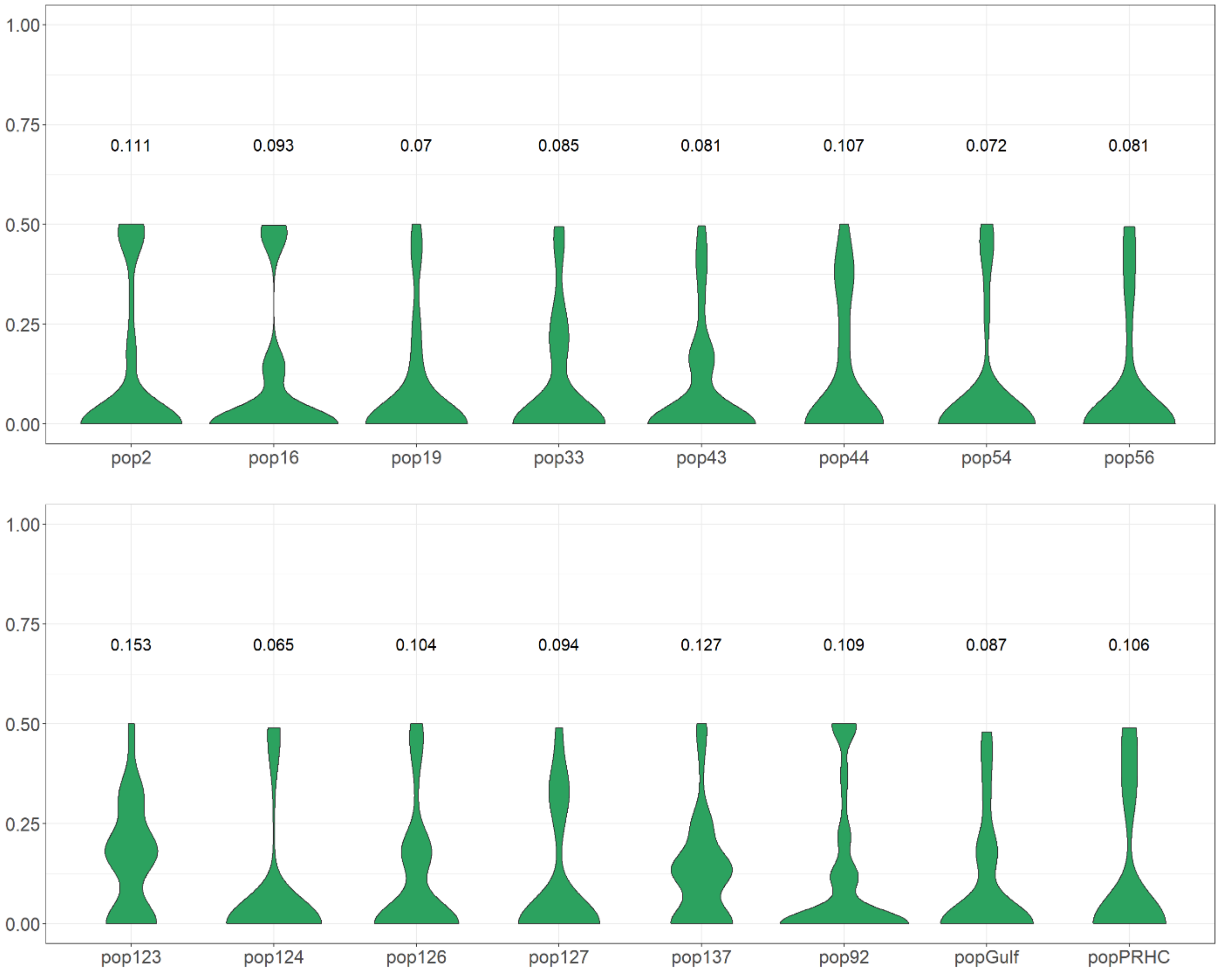


Supplemental Figure 1. Violin plots of expected heterozygosity of each population. Number on top of plots represent average values for each population.

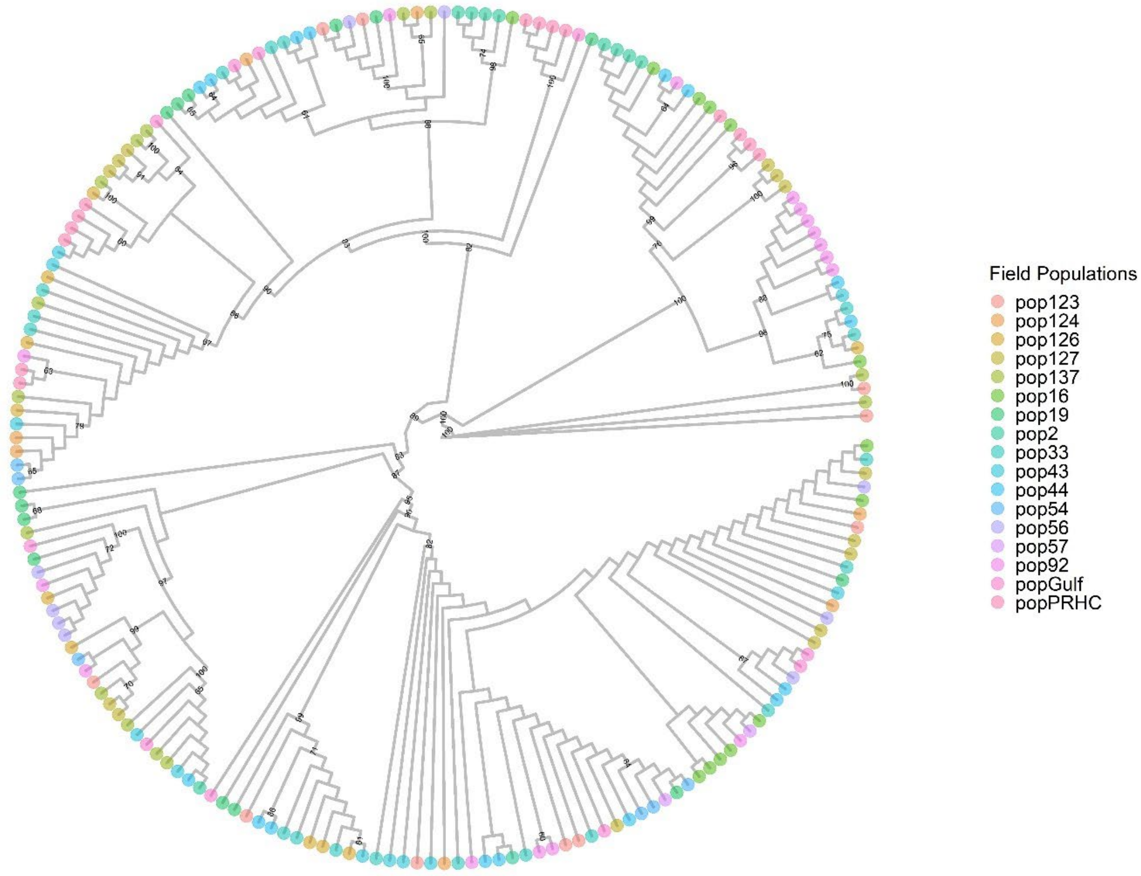


Supplemental Figure 2. Best-scoring maximum likelihood phylogenetic tree of *Lolium multiflorum* based on ACCase sequence. Numbers indicate bootstrapping values (values smaller than 60 were omitted). Tips were colored based on the population from which the haplotype originated.

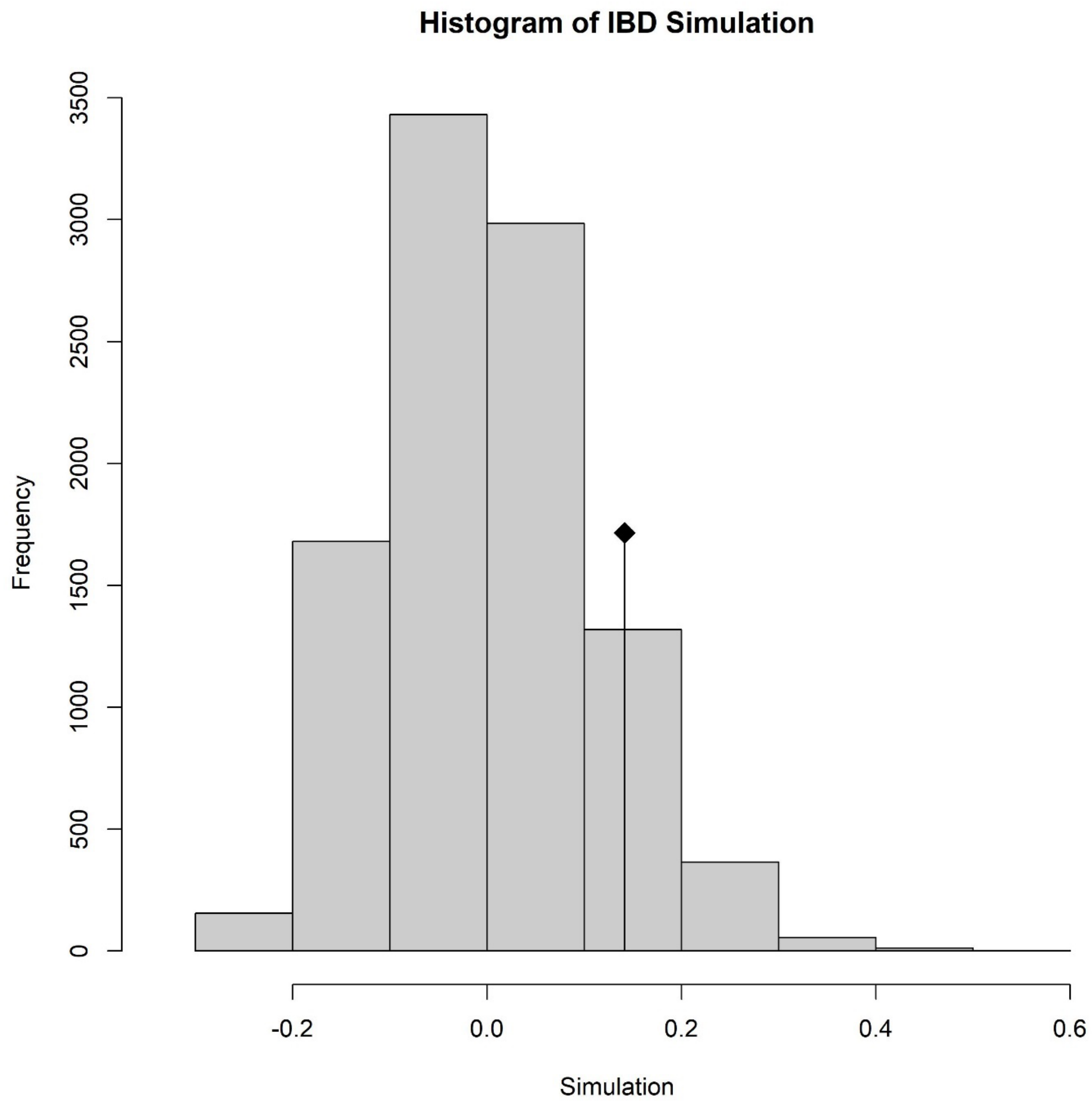


Supplemental Figure 3. Results of a Mantel test of correlation between genetic and geographic distance. Black diamond represents the value of the correlation based on our dataset. Bars represent a permutation with 10,000 replications to test the null hypothesis of absence of isolation-by-distance. A P value of 0.14 suggested there is no isolation-by-distance.

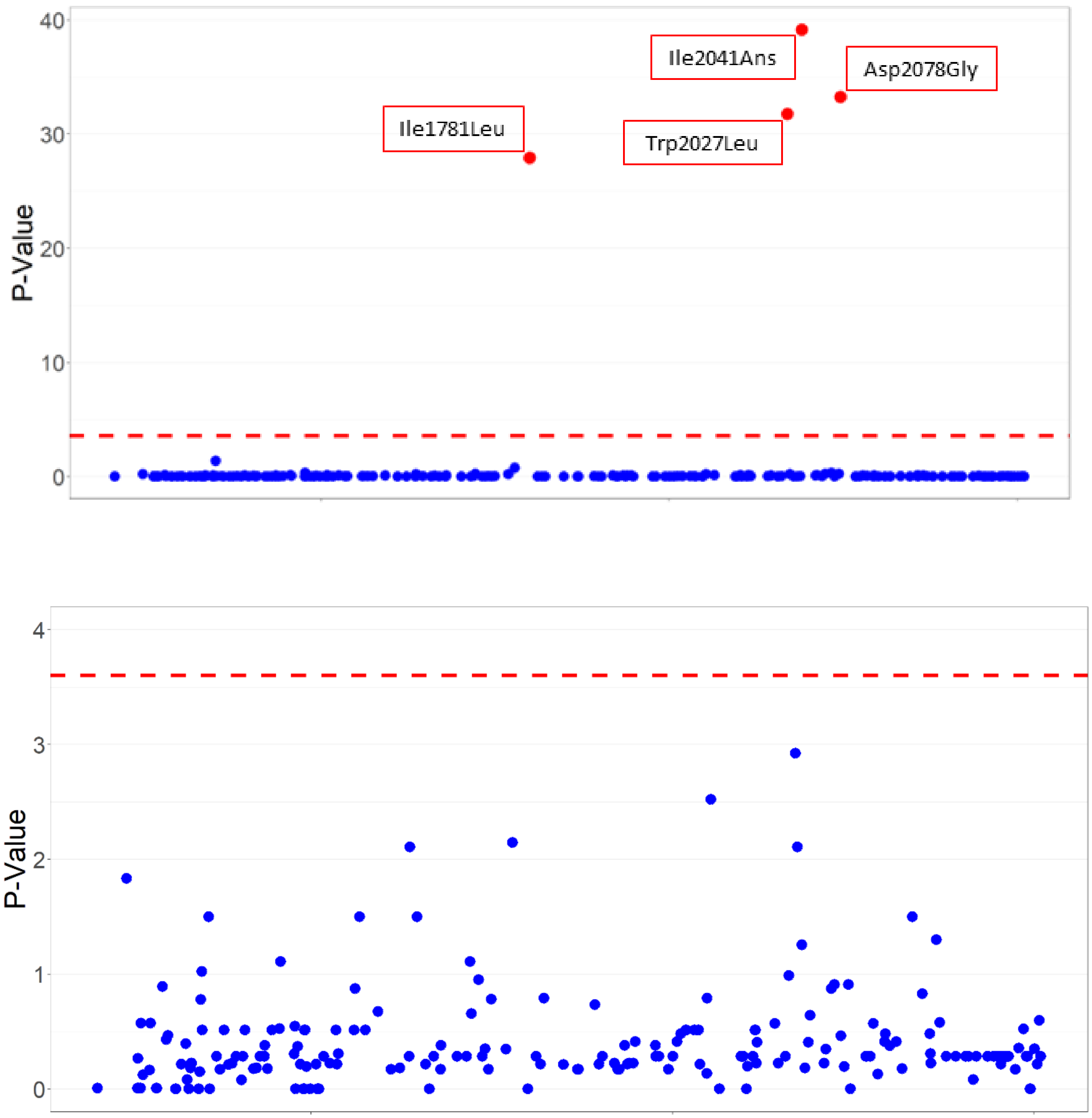


Supplemental Figure 4. Manhattan plots of association from analysis with amplicons from resistant individuals exhibiting SNPs known to confer ACCase herbicide resistance (Upper Panel, and amplicons with no known polymorphisms (Lower Panel). Red datapoints have P-value lower than adjusted cutoff. Dashed, red line represents cutoff P-value adjusted for false discovery rate.

Table S1. Geographical location and resistance phenotype.

| Population | Latitude | Longitude | Crop | Cross resistance[1] |
|---|---|---|---|---|
| pop2 | 45.55358 | -123.11542 | Orchardgrass | Clethodim, penoxaden, quizalofop |
| pop16 | 45.41847 | -122.97631 | Wheat | Clethodim, penoxaden, quizalofop |
| pop19 | 45.38315 | -122.88844 | Wheat | Clethodim, penoxaden, quizalofop |
| pop33 | 45.12463 | -123.18408 | Wheat | Quizalofop |
| pop43 | 45.05264 | -122.74122 | Tall fescue | Susceptible |
| pop44 | 45.04875 | -122.91118 | Wheat | Penoxaden, quizalofop |
| pop54 | 44.92392 | -122.70717 | Wheat | Clethodim, penoxaden, quizalofop |
| pop56 | 44.91136 | -122.75369 | Wheat | Clethodim, penoxaden, quizalofop |
| pop92 | 44.7255 | -123.00086 | Wheat | Clethodim, penoxaden, quizalofop |
| pop123 | 44.41946 | -123.27403 | Tall fescue | Penoxaden, quizalofop |
| pop124 | 44.4082 | -123.1399 | Wheat | Susceptible |
| pop126 | 44.40235 | -123.1242 | Tall fescue | Susceptible |
| pop127 | 44.39778 | -123.28228 | Tall fescue | Penoxaden, quizalofop |
| pop137 | 44.3503 | -123.13909 | Tall fescue | Susceptible |
| popPRHC[2] | - | - | Reference R | Penoxaden, quizalofop |
| popGulf[3] | - | - | Reference S | Susceptible |

[1]Resistance data is based on analysis of plant biomass. Populations that had a resistance index greater than 2 were classified as resistant. [2]Previously characterize ACCase inhibitor resistant population from Brunharo and Hanson (2018). [3]Known susceptible population from Brunharo and Streisfeld (2022).

Supplemental Table S2. Number of *Lolium multiflorum* individuals with and without amino acid substitutions in ACCase exhibiting resistance to at least one ACCase inhibitor.

| **Population** | **Individuals sequenced** | **Resistant individuals** | **SNPs found** | **Resistant without known SNPs** |
|---|---|---|---|---|
| pop2 | 5 | 4 | 2041, 2078 | 0 |
| pop16 | 7 | 6 | 2041, 2078 | 0 |
| pop19 | 4 | 3 | 1781, 2027, 2041, 2078 | 1 |
| pop33 | 10 | 6 | 2027 | 5 |
| pop43 | 6 | 1 | 2078 | 0 |
| pop44 | 8 | 7 | 2041 | 3 |
| pop54 | 4 | 4 | 1781, 2078 | 0 |
| pop56 | 5 | 4 | 2078 | 0 |
| pop92 | 8 | 7 | 2027, 2041, 2078 | 5 |
| pop123 | 5 | 4 | 2041 | 2 |
| pop124 | 4 | 0 | - | 0 |
| pop126 | 6 | 2 | 2027, 2078 | 0 |
| pop127 | 7 | 4 | 2041 | 1 |
| pop137 | 6 | 1 | 2041 | 0 |
| popPRHC | 7 | 4 | 1781, 2027 | 0 |
| popGulf | 5 | 0 | - | 0 |
| Total | 97 | 57 | - | 17 |

Supplemental Table 3. Dose-response regression estimates for clethodim, pinoxaden, and quizalofop based on biomass reduction compared to nontreated control.

| | **Clethodim** | | **Pinoxaden** | | **Quizalofop** | |
|---|---|---|---|---|---|---|
| Population | $GR_{50}$ | RI | $GR_{50}$ | RI | $GR_{50}$ | RI |
| pop2 | 203.0 ± 10.6 | 9.4 ± 0.7*** | 140.3 ± 7.9 | 11.1 ± 0.8*** | 136.6 ± 8.6 | 7.9 ± 0.6*** |
| pop16 | 114.3 ± 12.1 | 5.3 ± 0.3*** | 58.6 ± 5.4 | 4.6 ± 0.5*** | 222.4 ± 523.1 | 12.9 ± 30.4$^{ns}$ |
| pop19 | 283.8 ± 19.0 | 13.1 ± 1.1*** | 253.4 ± 15.8 | 19.9 ± 1.8*** | 628.8 ± 155.1 | 36.5 ± 9.3*** |
| pop33 | 25.8 ± 1.2 | 1.2 ± 0.0*** | 18.8 ± 0.6 | 1.5 ± 0.0*** | 35.5 ± 5.7 | 2.1 ± 0.4* |
| pop43 | 50.1 ± 3.3 | 2.3 ± 0.2*** | 17.7 ± 0.6 | 1.4 ± 0.0*** | 25.6 ± 1.3 | 1.5 ± 0.1*** |
| pop44 | 31.4 ± 1.7 | 1.4 ± 0.1*** | 116.1 ± 14.7 | 9.1 ± 1.3*** | 646.2 ± 71.7 | 37.5 ± 4.6*** |
| pop54 | 292.1 ± 25.4 | 13.5 ± 1.4*** | 213.5 ± 57.3 | 16.8 ±4.7** | 564.5 ± 125.0 | 32.8 ± 7.6*** |
| pop56 | 312.1 ± 32.7 | 14.4 ± 1.7*** | 171.4 ± 24.1 | 13.5 ± 2.1*** | 346.8 ± 38.1 | 20.1 ± 2.6*** |
| pop92 | 30.7 ± 2.1 | 1.4 ± 0.1*** | 190.9 ± 24.1 | 15.1 ± 2.3*** | 349.4 ± 25.9 | 20.3 ±1.9*** |
| pop123 | 43.8 ± 3.6 | 2.0 ± 0.2*** | 122.5 ± 10.1 | 9.7 ± 1.0*** | 22.9 ± 1.8 | 1.3 ± 0.1* |
| pop124 | 19.4 ± 1.7 | 0.9 ± 0.0$^{ns}$ | 13.8 ± 0.5 | 1.1 ± 0.0$^{ns}$ | 15.9 ± 0.7 | 0.9 ± 0.1$^{ns}$ |
| pop126 | 23.2 ± 1.2 | 1.0 ± 0.0$^{ns}$ | 11.1 ± 0.4 | 0.9 ± 0.0** | 19.5 ± 1.1 | 1.1 ± 0.1$^{ns}$ |
| pop127 | 46.6 ± 3.4 | 2.1 ± 0.2*** | 19.3 ± 2.6 | 1.5 ± 0.2* | 282.5 ± 38.4 | 16.4 ± 2.5*** |
| pop137 | 26.4 ± 1.9 | 1.2 ± 0.1* | 15.9 ± 0.7 | 1.3 ± 0.1*** | 18.1 ± 0.8 | 1.0 ± 0.1$^{ns}$ |
| popPRHC | 43.1 ± 3.4 | 2.0 ± 0.1*** | 22.8 ± 1.0 | 1.8 ± 0.1*** | 85.5 ± 12.3 | 4.9 ± 0.8*** |
| popGulf | 21.6 ± 1.2 | - | 22.8 ± 1.0 | - | 17.2 ± 1.0 | - |
| Total | | | | | | |

*P<0.05; **P<0.01; ***P<0.001. LL3 for all

Supplemental Table 4. Number of individuals exhibiting ACCase inhibitor resistance with known amino acid substitutions in ACCase.

| **Genotype** | **Quizalofop** | **Clethodim + quizalofop** | **Pinoxaden + quizalofop** | **Clethodim + pinoxaden + quizalofop** |
|---|---|---|---|---|
| I1781 | 2 | NA | 1 | 2 |
| W2027 | 3 | | | |
| I2041 | 1 | 2 | 4 | 8 |
| D2078 | NA | NA | NA | 10 |
| I1781 + I2041 | NA | NA | NA | 1 |
| I1781 + D2078 | NA | NA | NA | 1 |
| W2027 + I2041 | NA | 1 | 2 | NA |
| I2041 + D2078 | NA | NA | NA | 3 |
| I1781 + W2027 + I2041 + D2078 | NA | NA | 1 | NA |
